# Vitamin D deficiency induces erectile dysfunction: role of superoxide

**DOI:** 10.1101/2023.08.09.552595

**Authors:** Miguel A. Olivencia, Belen Climent, Bianca Barreira, Daniel Morales-Cano, Ana Sánchez, Argentina Fernández, Borja García-Gómez, Javier Romero-Otero, Claudia Rodríguez, Laura Moreno, Dolores Prieto, María Jesús Larriba, Angel Cogolludo, Javier Angulo, Francisco Perez-Vizcaino

## Abstract

Epidemiological studies suggest a relationship between vitamin D deficiency and erectile dysfunction (ED) but its causal relationship and the mechanism involved are unclear. Here we demonstrate that isolated corpora cavernosa (CC) from human donors with low vitamin D levels show reduced NO-dependent erectile function. This ED is also reproduced in vitamin D deficient rats and vitamin D receptor knockout mice *in vivo* and *ex vivo* and is associated with penile fibrosis. Vitamin D deficiency also blunts the response of CC to the phosphodiesterase 5 inhibitor sildenafil. CC from deficient rats show increased superoxide and their impaired erectile function is restored by superoxide scavengers. These results suggest that vitamin D deficiency induces ED via increased superoxide production.

## INTRODUCTION

Erectile dysfunction (ED) is characterized by the inability to achieve or maintain an erection of the penis during sexual activity ^1^. It is a highly prevalent condition, affecting 30% of European (40-79 years) and 52% of US (40-70 years) males ^2^. Moreover, it adversely impacts the quality of life and is considered a sentinel marker for poor general health ^3^. In fact, ED has independent predictive value for future myocardial infarction and stroke ^4,5^. Obesity, metabolic diseases, and diabetes are well-recognized risk factors for sexual dysfunction ^5,6^. Impaired bioactivity of NO released by nerve and endothelial cells in the corpora cavernosa (CC) of the penis is a major pathogenic mechanism in ED ^5-7^. Contemporary treatment algorithms for ED involve the use of phosphodiesterase 5 inhibitors (PDE5i) as first choice agents to potentiate the NO/cyclic GMP pathway ^5,8^. It is estimated that 30% of patients with ED are non-responders to PDE5i ^9^ and intracavernosal injection of prostaglandins or other vasoactive agents remains as an alternative therapy ^5^.

Vitamin D, mainly synthesized in skin after UV exposure, has physiological functions beyond calcium and phosphorus homeostasis, including the regulation of cellular growth, intracellular metabolism, or innate and adaptive immunity ^10^. Vitamin D status is an important health issue since more than the half of the world population shows vitamin D deficiency ^10,11^ (25-hydroxyvitamin D plasma levels below 20ng/mL ^12,13^). Epidemiological studies have reported a higher prevalence of vitamin D deficiency in ED patients, an association between the severity of ED with 25-hydroxyvitamin D plasma levels ^14-16^ and an improved response to PDE5i after vitamin D replacement ^17^. However, the cause-effect relationship is unclear, and the potential mechanisms are unknown.

Herein, we show for the first time that isolated human CC from organ donors with vitamin D deficiency exhibit impaired erectile function and that this is reproduced *ex vivo* and *in vivo* in rats with vitamin D deficiency, and in vitamin D receptor knockout mice. Our data indicates that this is due to increased radical superoxide.

## RESULTS

### Vitamin D levels are associated with ED *in vitro* in human donors

The aetiologies of ED include vascular, hormonal, neurologic, and/or psychological dysfunctions which are often interlinked ^5^. We interrogated whether the relationship between ED and vitamin D deficiency reported in epidemiological studies could be also observed ex *vivo*, i.e., in the absence of hormonal or psychological influences. Therefore, we analysed 25-hydroxyvitamin D levels in the plasma from organ donors and compared them with the relaxant responses induced by electrical field stimulation (EFS) in their isolated human CC. Donor characteristics are shown in Supplemental Table 1. EFS induces the release of NO from nitrergic nerves, inducing arterial vasodilation and relaxation of the trabecular smooth muscle cells of the CC, causing an increase in intracavernosal pressure and, hence, penile engorgement. Endothelium- and NO-dependent vasodilation of penile arteries is also an important contributor to penile erection. Thus, EFS-induced relaxation represents an *ex vivo* surrogate test for erectile function while acetylcholine-(ACh) induced vasodilation represents a standard test to analyse endothelial function. In agreement with the high prevalence of both ED and vitamin D deficiency, many of the donors show moderately or severely reduced 25-hydroxyvitamin D levels and/or reduced responses to EFS in CC (Fig. 1). Both parameters were normally distributed in the cohort. Interestingly, we found a significant direct correlation between both parameters, and, therefore, erectile function was markedly different in patients with 25-hydroxyvitamin D levels above vs. below the median value (14.2 ng/ml) of the cohort (Fig. 1A and 1B). A similar correlation was found between 25-hydroxyvitamin D levels and the endothelial response stimulated with Ach in human penile resistance artery (HPRA). Again, the median 25-hydroxyvitamin D levels clearly discriminated between those with high and low endothelial-dependent response (Fig. 1C). However, testosterone in plasma did not correlate with 25-hydroxyvitamin D levels (supplemental Fig. 1A). Thus, even when the sample size was small, the results strongly suggest that the association of vitamin D deficiency with poor erectile function remains in isolated human CC.

**Fig. 1.**
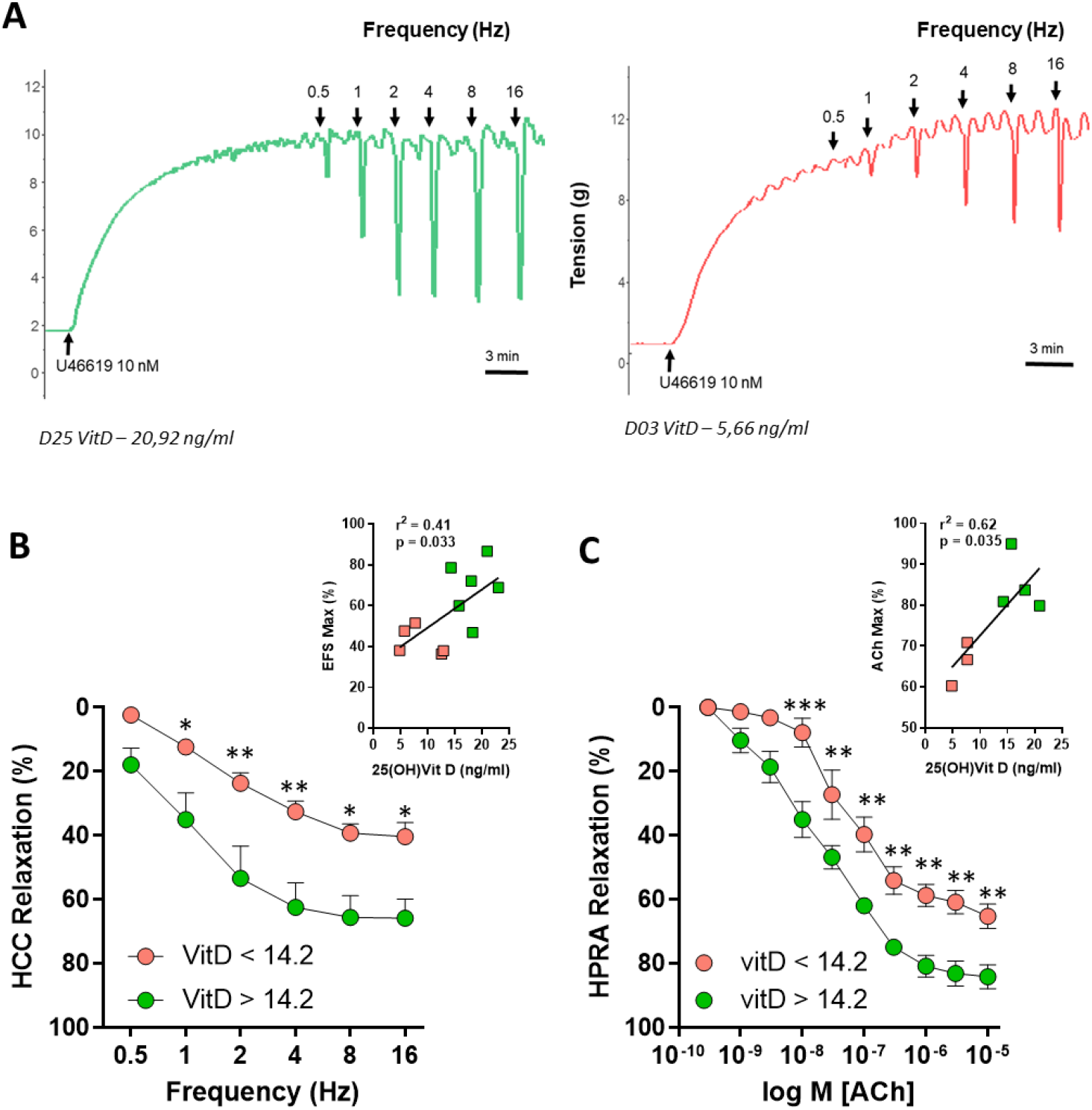
25-hydroxyvitamin D plasma levels correlate with human corpus cavernosum (CC) and human penile resistance artery (PRA) function ex vivo. 25-hydroxyvitamin D plasma levels were measured in donors and HCC were divided into two groups, those from patients with 25-hydroxyvitamin D above or those below the median value (14.2 ng/mL). A) Original recordings of human CC stimulated by EFS from a patient with low (left) or normal (right) 25-hydroxyvitamin D levels. B) Relaxant responses of human CC induced by EFS in the two groups and its correlation with 25-hydroxyvitamin D plasma levels. C) Relaxant responses of human PRA induced by ACh and its correlation with vitamin D plasma levels. Results are means ± standard error of the mean (sem). *, ** P<0.05 and P<0.01, respectively, vs low 25-hydroxyvitamin D by two-way (deficit x frequency or deficit x concentration) ANOVA test followed by a Sidak’s multiple comparisons test. A Pearson correlation coefficient was calculated for the relationship between 25-hydroxyvitamin D plasma levels and maximal responses.

### Vitamin D deficiency induces *in vivo* and *ex vivo* ED and penile fibrosis

To analyse the cause-effect relationship, in parallel with the pulmonary study of a previous project ^18^, we evaluated the erectile function in wild type Sprague rats exposed to a vitamin D-free diet for 5 months in comparison with a standard diet. These preliminary results indicated that a decrease in 25-hydroxyvitamin D levels ^18^ results in a significant reduction in EFS-induced relaxation (Supplemental Fig. 2A). In addition to neuronal-derived NO, NO released from the endothelium of sinusoids and blood vessels may also be involved in the penile erection. Vitamin D deficiency also induced a significant decrease in the endothelium NO-dependent relaxation induced by ACh (Supplemental Fig. 2B). Moreover, the relaxation induced by the PDE5i sildenafil, which potentiates endogenous NO, was also reduced (Supplemental Fig. 2C).

However, the relaxations to the soluble guanylyl cyclase (sGC) stimulator riociguat did not change (Supplemental Fig. 2D). The relaxation induced by EFS was almost entirely inhibited by the NO synthesis inhibitor L-Nitro-Arg and potentiated by sildenafil in both groups (Supplemental Fig. 3). In addition, the relaxation to ACh (Supplemental Fig. 4A) and sildenafil (Supplemental Fig. 4B) in the dorsal penile arteries was also reduced in animals with vitamin D deficiency.

We designed a new set of experiments with Wistar rats exposed for 5 months to a vitamin D-free diet to analyse the erectile function *in vivo* and to gain insights into the mechanisms involved in ED. This intervention reduced significantly the 25-hydroxyvitamin D plasma levels but did not affect those of testosterone (Supplemental Fig. 5). Notably, the increase in intracavernosal pressure induced by electrical stimulation of the cavernosal nerve in anaesthetized rats was markedly decreased by vitamin D deficiency (Fig. 2A and 2B). In addition, CC strips isolated from vitamin D deficient rats also showed reduced relaxation induced by EFS (Fig. 2C) and by the PDE5i sildenafil (Fig. 2D). Therefore, vitamin D deficiency induced erectile dysfunction *ex vivo*, confirming the pilot study, and *in vivo*, which was associated with reduced response to sildenafil, the first-choice therapeutic drug for ED.

**Fig. 2.**
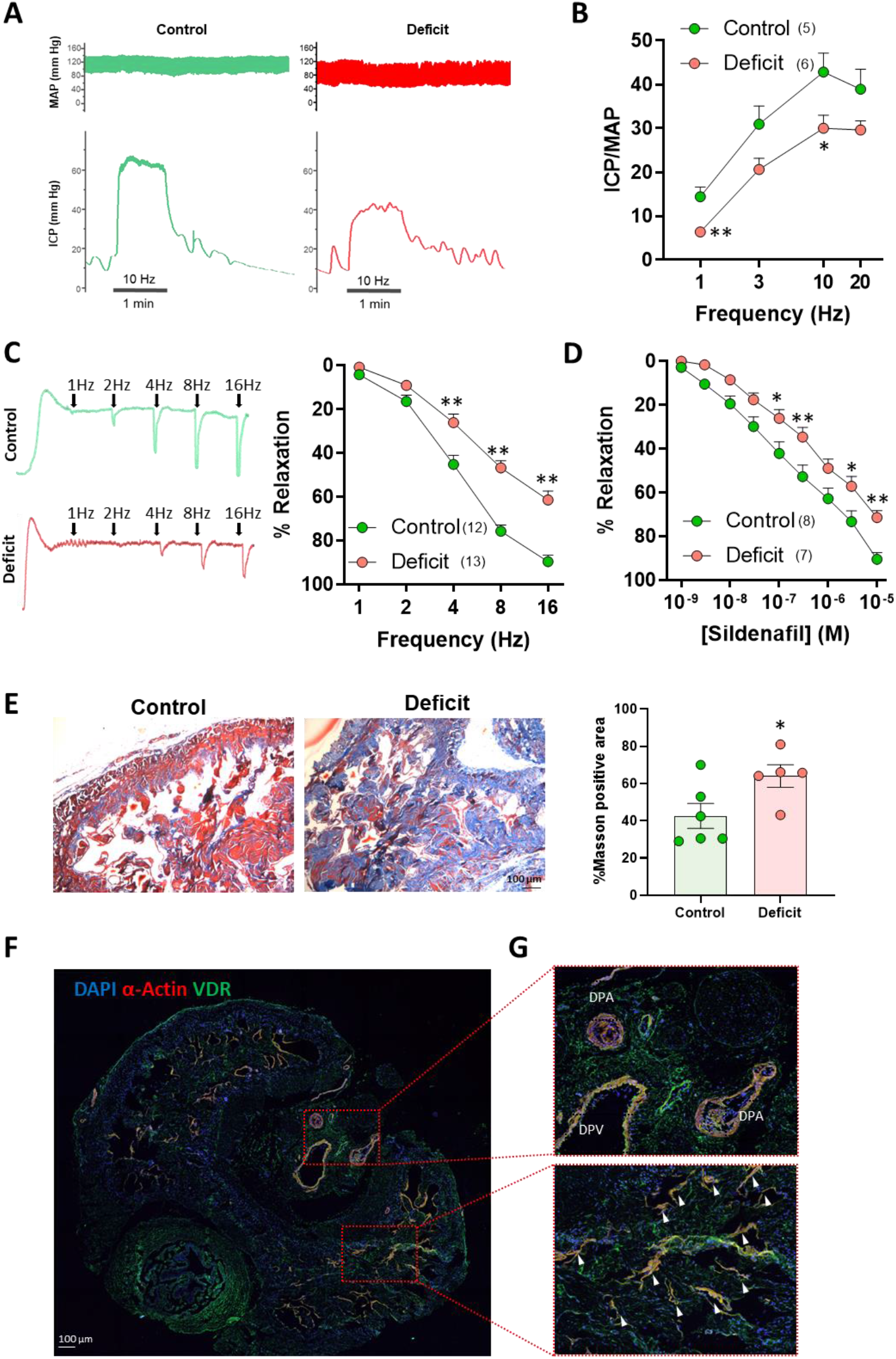
Vitamin D deficiency induces ED in rats in vivo and ex vivo. A) Original traces of the mean arterial pressure (MAP) and intracavernosal pressure (ICP) recordings after electrical stimulation of the cavernous nerve in anaesthetized rats at 10Hz of frequency. B) Averaged increases in ICP normalized with MAP. C and D) Effects of vitamin D deficiency on the relaxant responses of CC induced by C) EFS including original traces and D) sildenafil. E) Representative images of cross-sections of penises stained with Masson trichrome. Extracellular collagen deposition was quantified as the percentage of blue over the total area, 4-7 images per animal were used. Results are means ± sem. F) VDR localization by immunofluorescence labelling and confocal imaging in penis rat slices. VDR expression is shown in green, α-actin in red and nuclei are shown in blue (DAPI). DPA = Dorsal penile artery; DPV = Dorsal penile vein. Arrows indicate colocalization of VDR and α-actin. *,** P<0.05 and P<0.01, respectively, vs control using t-student test (for panel E) or otherwise using two-way (deficit x frequency or deficit x concentration) ANOVA test followed by a Sidak’s multiple comparisons test.

Penile cavernous fibrosis is considered an important factor leading to ED. Therefore, we analysed the collagen deposition in paraffin penis sections by Masson trichrome staining, as a marker of fibrosis. We found that vitamin D deficiency significantly increased the blue stained area indicating excess deposition of collagen in the CC (Fig. 2E).

The active form of vitamin D, calcitriol, activates vitamin D receptor (VDR), a member of the nuclear receptor superfamily of transcription factors that regulate gene expression. To confirm that VDR is present in the rat penis, its expression was analysed in sections of penises from Wistar rats by immunohistochemistry. Fig. 2F shows a positive staining for VDR in cavernosal smooth muscle cells, the dorsal penile arteries and the dorsal penile vein where it colocalized with smooth muscle α-actin.

### *Vdr* knock-out mice exhibit erectile dysfunction *ex vivo* and *in vivo*

Then, we analysed whether the ED induced by vitamin D deficiency could be also mimicked by genetic deletion of its receptor. *In vivo* and *ex vivo* erectile function was explored in *Vdr* knockout (*Vdr*^*-/-*^) and WT mice. As shown in the original recordings (Fig. 3A) and in the averaged data (Fig. 3B), the increase in ICP in response to electrical stimulation was lower at all electrical frequencies in *Vdr*^-/-^ mice compared to WT, confirming the erectile dysfunction *in vivo*. Furthermore, CC isolated from *Vdr*^-/-^ mice showed decreased relaxation induced by EFS (Fig. 3C) and by the PDE5i sildenafil (Fig. 3D) also corroborating the ED *ex vivo*. However, as in vitamin D deficient rats, there were no changes in the response to the sGC stimulator riociguat in *Vdr*^*-/-*^ mice (Fig. 3E).

**Fig. 3.**
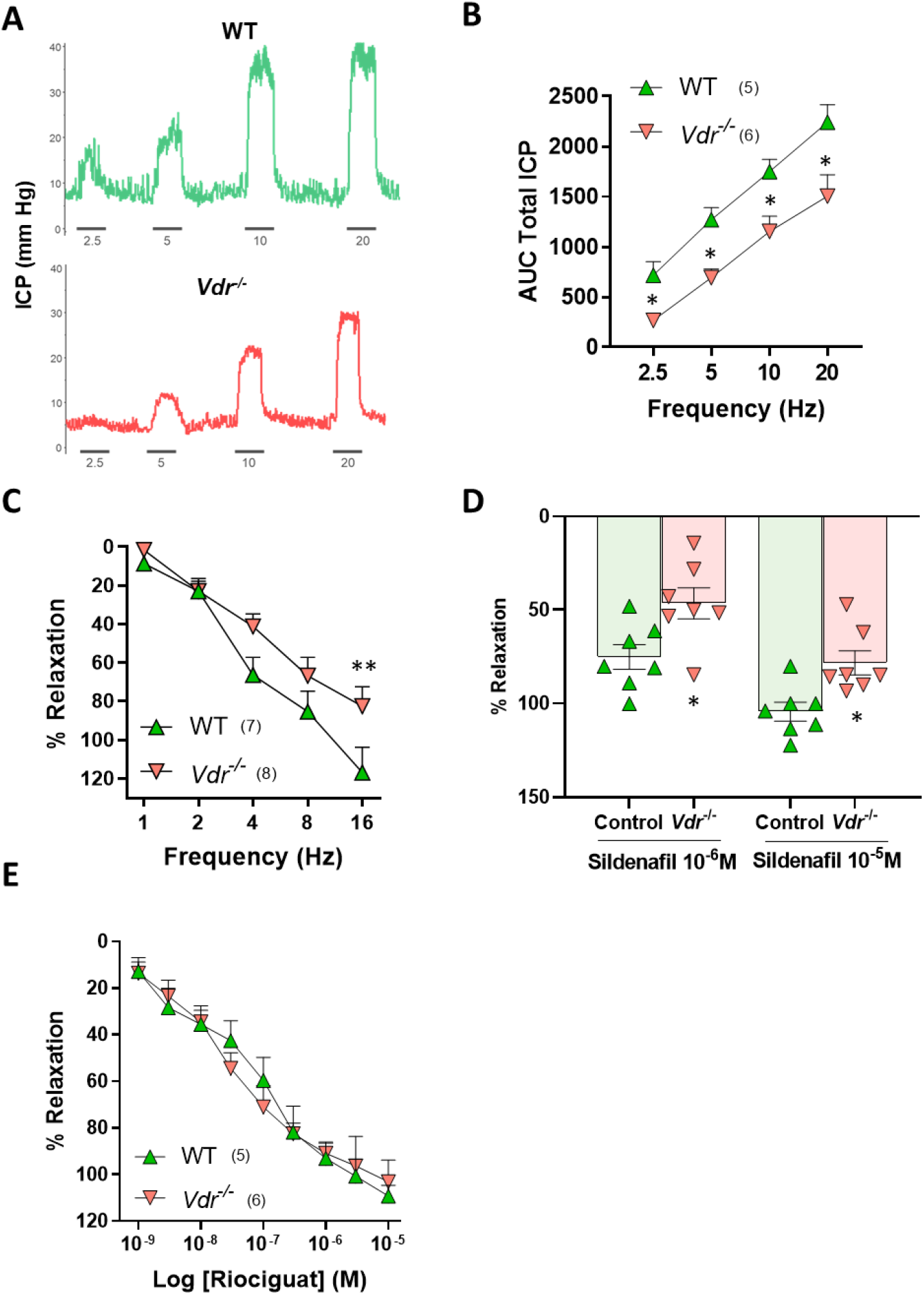
Vitamin D receptor knockout induces ED in mice ex vivo and in vivo. A) Original traces of intracavernosal pressure recording after electrical stimulation of the cavernous nerve in anaesthetized WT and Vdr-/-mice with increasing frequencies. B) Averaged increases in pressure. C, D, E) Effects of Vdr deletion on the relaxant responses of CC induced by C) EFS, D) sildenafil, E) Riociguat. A full frequency- or concentration-response curve was performed for EFS and riociguat but only two concentrations of sildenafil were tested because the responses were slower to this drug. Results are means ± sem. *,** indicates P<0.05 and P<0.01, respectively, vs WT using two-way (deficit x frequency or deficit x concentration) ANOVA test followed by a Sidak’s multiple comparisons test.

### Excess superoxide mediates vitamin D deficiency- and *Vdr* deletion-induced ED

The reduced responses to EFS and sildenafil in vitamin D deficient rats and *Vdr*^*-/-*^ mice, which are dependent of endogenous NO, and the unaffected response to riociguat, which mimics NO but does not require endogenous NO, gave us a clue of a possible mechanism: a reduced bioavailability of NO due to excess levels of superoxide anion, which is a well-known mechanism involved in ED ^19^. Thus, we measured superoxide production in human penile sections by DHE fluorescence and compared these data with the *ex vivo* function of the CC and the 25-hydroxivitamin D levels in each donor. As expected, donor tissue sections with higher intensity of DHE fluorescence exhibited reduced maximal relaxant response to EFS (Fig. 4A), i.e., the higher the superoxide, the worse erectile function. Interestingly, DHE signal also correlated with the levels of 25-hydroxyvitamin D (Fig. 4B).

**Fig. 4.**
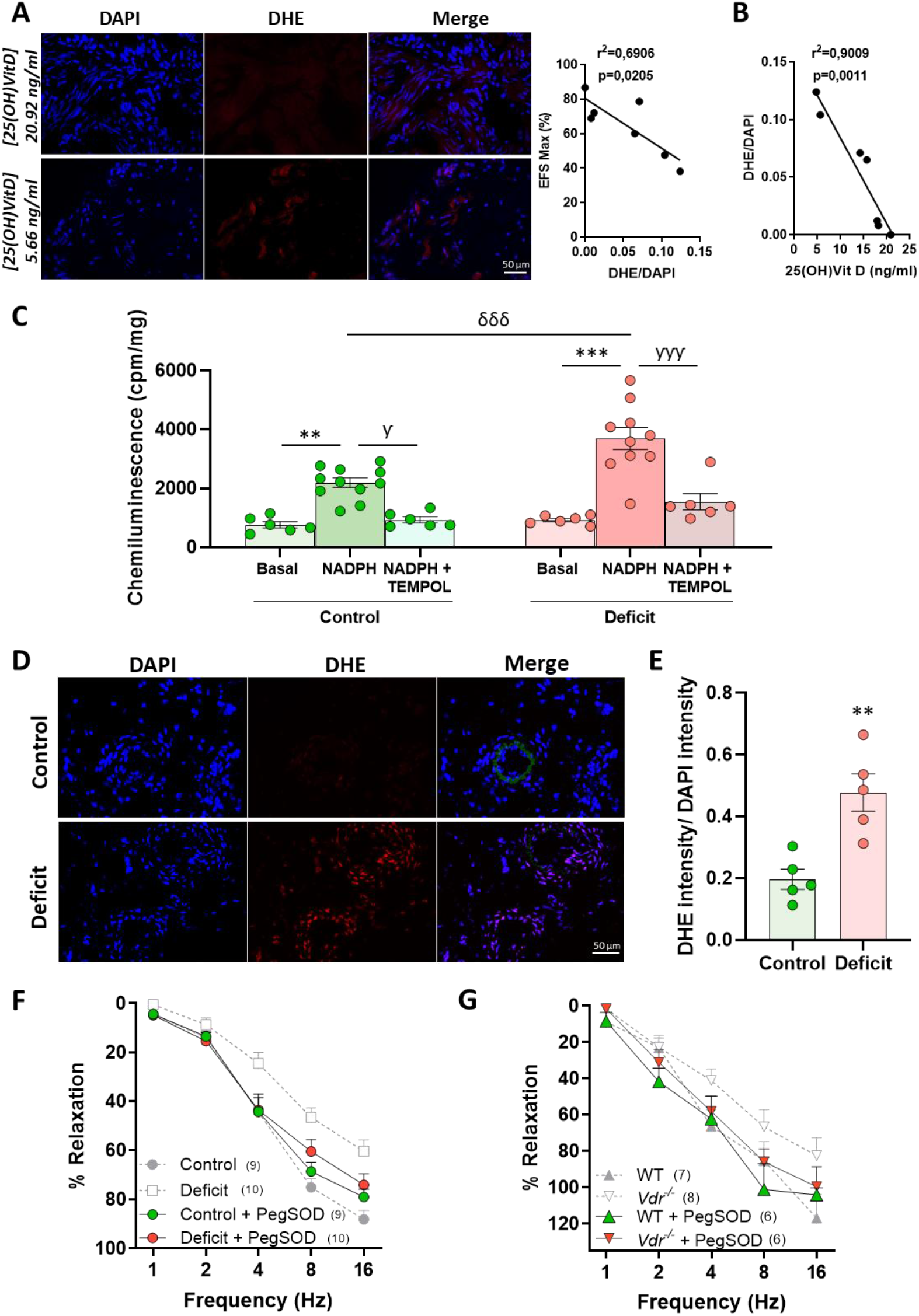
Vitamin D deficiency- and Vdr deletion-induced erectile dysfunction is mediated by increased superoxide. A) Dihydroethidium (DHE) staining in the CC of a donor with high (left) and low (right) 25-hydroxivitamin D. B) Correlation between DHE intensity normalised by DAPI (DHE/DAPI) and the maximal effect of electrical field stimulation (EFS Max) (left) and between 25-hydroxyvitamin D plasma levels and DHE/DAPI (right) in CC of human donors. C) Levels of superoxide (O^2•-^) measured by lucigenin chemiluminescence in rat CC strips in the absence (basal) or in the presence of NAPDH or NADPH plus TEMPOL. D) Rat penis slices showing the blue fluorescence of the nuclear stain DAPI (left), the red fluorescence produced when DHE is oxidized to ethidium by superoxide (middle), and the merged images (right). E) Values of DHE/DAPI. 4-10 images were taken from each animal. F, G) Effects of the relaxant responses of CC induced by EFS in presence or absence of the antioxidant PegSOD from F) vitamin D deficient rats, G) Vdr knockout mice. The results from figures 2C and 3C are superimposed in light grey and discontinuous lines just for reference. Results are means ± sem. ** indicates P<0.01, ***P<0.001 vs basal or control; ƴP<0.05, ƴƴƴP<0.001 vs NADPH, δδδP<0.01 vs control NADPH using one way ANOVA test followed by a Bonferroni post hoc test or student’s t test in panel C.

To analyse the causal relationship between 25-hydroxyvitamin D and superoxide, we measured superoxide levels by both lucigenin luminescence and DHE staining in CC isolated from control and vitamin D deficient rats. In CC strips from vitamin D deficient rats, the basal lucigenin signal was not significantly different from the control strips (Fig. 4C), but the NADPH-induced increase was significantly higher in CC from vitamin D deficient rats. In both experimental groups the antioxidant tempol prevented the NADPH-induced signal, indicating that it was specific for superoxide production. Similarly, the penises from vitamin D deficient rats showed marked red DHE nuclear staining (Fig. 4D,E), again indicating high superoxide levels, compared to controls.

To explore whether increased superoxide generation was involved in the ED, we analysed the response to EFS of CC strips from control and vitamin D deficient rats and from *Vdr*^*-/-*^ and WT mice incubated with pegylated superoxide dismutase (PegSOD). This antioxidant reverted the *in vitro* ED induced by both vitamin D deficiency in rats (Fig. 4F) and by genetic ablation of the receptor in mice (Fig. 4G). Therefore, these data strongly suggest that vitamin D deficiency- and VDR ablation-induced ED was mediated by increased superoxide production.

## DISCUSSION

Previous epidemiological studies have associated vitamin D status with erectile function. Moreover, the 25-hydroxyvitamin D plasma level has been proposed as an independent risk factor for ED ^14-16^ and was also related with the severity of ED. Here, we report for the first time that vitamin D levels correlate with EFS-induced relaxation in isolated CC and with endothelial function in isolated resistance penile arteries from human donors. This study also demonstrates, in rats and mice, a causal relationship between exposure to vitamin D-free diet or *Vdr* knockout and ED both *ex vivo* and *in vivo*. Moreover, we suggest that increased superoxide is involved in vitamin D deficiency-mediated ED.

Vitamin D deficiency may potentially impact most aetiologies of ED. In this regard, low levels of 25-hydroxyvitamin D are linked to depressive disorders, a possible source of psychogenic ED ^20-22^. Optimal testosterone levels, essential for correct erectile function, have also been correlated with adequate 25-hydroxyvitamin D levels ^23^. Our results indicate that the cause of the failure in penile erection may be intrinsic to the CC and that neither human nor rat testosterone levels are influenced by 25-hydroxyvitamin D levels. Consequently, vitamin D deficiency induces ED of vascular origin, although psychogenic and hormonal aetiologies cannot be completely ruled out. Nevertheless, our studies in human CC must be interpreted with caution because the relatively low small number of samples and because the association might not be causal.

In rats exposed to vitamin D-free or to the standard diet, 25-hydroxyvitamin D levels are comparable to those found in the two categories of human subjects studied, above and below the median level of 25-hydroxyvitamin D levels, respectively. Therefore, vitamin D-free diet is a suitable model to study ED. In addition, a recent study ^24^ reports that supplementation with vitamin D3 above the standard levels improves erectile function in a rat model of cavernous nerve injury.

When the active form of vitamin D, calcitriol, binds VDR, it acts as a transcription factor and interacts with the VDRE in the promoter region of target genes regulating their expression ^25,26^. In our study, we observed ubiquitous VDR expression in the penis, and, in particular, in the smooth muscle cells of the vessels and the CC. Interestingly, our results indicate that the absence of VDR causes erectile dysfunction *in vivo* and *ex vivo*.

Penile cavernous fibrosis is considered an important factor leading to ED. The pro-fibrotic factors TGF-β and its downstream signalling partners p-SMAD2 and p-SMAD3 play a pathophysiological role in neurogenic ^27^ and vasculogenic ^28^ models of ED and activated TGF-β signalling has also been observed in smooth muscle and endothelial cells from patients with ED ^29^. We also find fibrosis with increased collagen deposition in the CC from vitamin D deficient rats. This is consistent with the well-known anti-fibrotic effect of vitamin D in other organs such as the liver ^30^, lung ^31^ or kidney ^32^ which is widely attributed to its double, VDR-dependent and -independent, inhibitory effect on TGF-β signalling ^33^.

Impairment of the NO pathway is recognized as a major pathogenetic factor in animal models and patients with ED ^34-36^. As expected, the relaxation induced by EFS in both experimental groups is NO-dependent since the NO synthesis inhibitor L-NAME suppressed it. Likewise, sildenafil, that inhibits the degradation of cyclic GMP, the downstream signal of NO, potentiated EFS in both groups. Moreover, vitamin D deficiency significantly reduced the responses to acetylcholine and sildenafil in both the CC and the penile dorsal artery. Accordingly, the relaxation of the sGC stimulator riociguat in CC, which does not require endogenous NO, was unaffected by vitamin D deficiency or *Vdr* ablation. This is consistent with a reduced bioavailability of NO.

The finding that vitamin D deficiency induces a reduced response to PDE5i in both the CC and the penile arteries may have additional clinical implications. For unclear reasons, many patients with ED are non-responders to PDE5i ^9^. Our results suggest that vitamin D deficiency, may contribute to the lack of response to this first-line treatment of ED. Likewise, addition of vitamin D to the PDE5i tadalafil increased the erectile function in patients with ED ^17^. Similarly, vitamin D deficiency reduces the effect of sildenafil in rat pulmonary arteries and is a good predictor of poor therapeutic response to PDE5i in patients with pulmonary arterial hypertension ^37^.

Oxidative stress is an important factor in the development of ED ^19^ and NADPH oxidase has been reported to be the main generator of superoxide in the CC in vascular ED ^19,38,39^. Accordingly, our results of increased NADPH-stimulated lucigenin luminescence and dihydroethidium fluorescence indicated an increase in superoxide penile production after chronic vitamin D deficiency. The functional experiments with pegSOD confirmed the causative role of superoxide in vitamin D deficiency- or *Vdr* knockout-induced ED. Interestingly, preceding this study, low vitamin D levels have been extensively related to oxidative stress-induced vascular dysfunction ^40,41^. In this regard, vitamin D has been described to reduce oxidative stress through NADPH oxidase downregulation ^41,42^. Additionally, the protective role against oxidative stress of vitamin D has been attributed to increased levels of glutathione (GSH) ^43-45^ or increased expression of the nuclear factor erythroid 2-related factor 2 (Nrf2) ^46-48^. Nevertheless, the underlying mechanism whereby vitamin D deficiency induces ED mediated by increased superoxide production is unknown.

Systemic endothelial dysfunction and ED essentially arise from a failure of the NO/cyclic GMP pathway, primarily caused by an excess of superoxide-driven NO inactivation, and both conditions are early prognostic markers for future cardiovascular events. Endothelial dysfunction is silent and has been widely associated with vitamin D deficiency ^49,50^. Our study provides a causal relationship and the mechanistic basis for the association of ED with vitamin D deficiency.

In conclusion, we demonstrate for the first time that vitamin D deficiency or *Vdr* knockout induce erectile dysfunction in rodents. We suggest that this is mediated by increased CC superoxide production. We further suggest that vitamin D deficiency is an aetiological factor for vascular ED and for the therapeutic failure of PDE5i. Moreover, our results raise the possibility that restoring the vitamin D status in patients with vitamin D deficiency and ED would improve not only its calcium metabolism and bone health but also its sexual performance and/or the efficacy of the treatment for ED.

## MATERIALS AND METHODS

### Human tissues

Cavernosal specimens and blood samples were obtained from 12 organ donors at the time of organ collection for transplantation in the Hospital Universitario Doce de Octubre, Madrid, Spain. In addition to provide consent for organ transplantation, relatives provided informed consent for tissue procurement specifically for research. The project protocol was approved by the Ethics Committees of the Hospital Universitario Doce de Octubre, Madrid, Spain (Ethics Approval procedure 16/045) and the Hospital Universitario Ramón y Cajal, Madrid, Spain (Ethics Approval procedure 363-15). Subjects were only excluded when suffering any infectious disease. Tissues were maintained in sterilized M-400 solution (composition per 100 ml: mannitol, 4.19 g; KH_2_PO_4_, 0.205 g; K_2_HPO_4_·H_2_O, 0.97 g; KCl, 0.112 g; NaHCO_3_, 0.084 g; pH 7.4, at 4-6°C). Time elapsed between harvesting of cavernosal specimens from organ donors to their functional evaluation ranged between 16 and 24 hours. Within this time range, tissues are totally viable and are adequate for functional evaluation ^51^. Plasma was obtained by centrifugation at 4ºC of blood collected in EDTA-containing tubes.

### Animal model

Experimental procedures were approved by the Animal Welfare Body of Universidad Complutense and Consejo Superior de Investigaciones Científicas and the regional committee of Comunidad de Madrid (Ref. PROEX-016/019 and PROEX-396.1/21) in accordance with the guidelines on the ethical use of animals from the European Community Council Directive of September 22, 2010 (2010/63/EU). All investigators understand the ethical principles. All animals were kept under standard conditions of temperature 22±1°C and 12:12 hour dark/light cycle with free access to food and water.

A pilot study was performed in CC from wild-type male Sprague-Dawley (SD) rats with/without vitamin D-free diet from a previous protocol ^18^. In addition, another batch of two-month old male Wistars rats was randomly allocated into two groups: rats fed with a standard diet (Teklad Global 18% Protein Rodent Diet, Envigo, n=20) or vitamin D-free (Teklad Custom Diet TD.120008, Envigo, n=20) diet during 5 months. *Ex vivo* incubation experiments were performed with corpora cavernosa obtained from control male Wistar rats (n = 14).

The *Vdr* knockout (*Vdr*^-/-^, n=14) and wild-type (WT, n=14) mice were obtained by crossing heterozygous mice (Vdr^+/-^) ^52^ and genotyped as described (protocol 22517 in http://www.jax.org) and used at 4 months of age. *Vdr*^-/-^ were fed a g-irradiated diet (TD96348, Teklad, Madison, WI) containing 2% calcium, 1.25% phosphorus, and 20% lactose ^53^.

### Functional evaluation of CC and penile resistance arteries

All functional experiments were performed in Krebs-Henseleit solution at 37° C, pH 7.4 and bubbled with 95% O_2_ and 5% CO_2_. Strips of human CC tissue from eleven patients were mounted in organ chambers and stretched to optimal isometric tension as described ^51^. Neurogenic relaxations to EFS (0.5 to 16 Hz) were obtained in CC strips contracted with U46619 (10-30 nM). Penile small helicine arterial rings (lumen diameter 300-500 μm, 1.7-2.0 mm long), were dissected from seven CC specimens and mounted on microvascular wire myographs for isometric tension recordings as described ^54^. The arteries were contracted with 1-3 μM noradrenaline and the relaxation was evaluated by cumulative addition of acetylcholine (ACh).

CC strips from rats and mice were suspended between two electrodes allocated in a wire myograph, stretched to 3 mN and incubated with guanethidine (10 μM) and atropine (1 μM) to block adrenergic neurotransmission and muscarinic receptors, respectively. The preparations were contracted with phenylephrine (0.3-1 μM). Relaxations were induced by EFS stimulation or by vasoactive agents ^55^. Rat dorsal penile arterial segments (2 mm long) were mounted in a wire myograph and stretched to give an equivalent transmural pressure of 100 mmHg. Arteries were contracted with phenylephrine (1–3 μM) and relaxations were induced by ACh or sildenafil. All drugs were incubated for 10 min except PEGSOD for 30 minutes. Relaxations were expressed as a percentage of the reduction in vasoconstrictor-induced contraction.

### Intracavernosal pressure recording

Rats and mice were anaesthetized with ketamine (60 mg·kg^-1^) and diazepam (4 mg·kg^-1^). The right cavernous nerve was dissected and intracavernosal pressure (ICP) was recorded by insertion a 25-gauge needle into the right crus. Electrical stimulation was applied by a platinum bipolar hook electrode connected to a stimulator and frequency-response curves were performed by applying stimulation at increasing frequencies at 3 min intervals ^34,55^. Left carotid artery was catheterized in rats for constant blood pressure measurement.

### Histology

Tissue samples from the penis were either fixed in 4% paraformaldehyde in 0.1 M phosphate-buffered saline (PBS) and embedded in paraffin, or cryoprotected in 30% sucrose in PBS, immersed in OCT, snap frozen in liquid nitrogen, and stored at −80°C. Paraffin transversal penis sections were stained with Mason trichrome techniques and examined by light microscopy. Quantification of blue staining was performed with Image-Pro plus 2D Image Analysis Software in the whole section.

For immunostaining, OCT-embedded penis sections were permeabilized in PBS with 0.3% Triton X-100 at room temperature for 10 min following by 1h incubation with blocking buffer containing 10% normal goat serum (Life Technologies, 16210-072) and 0.1% Tween20 in PBS at room temperature. Then, slices were incubated with primary antibodies overnight at 4°C. The following immune fluorescence staining were performed: anti-VDR (1:50, sc-13133, mouse monoclonal, Santa Cruz, Dallas, USA) followed by Alexa594-conjugated anti-mouse (1:200, A11032, Thermo Fisher Scientific, Massachusetts, USA) and anti-α-smooth muscle actin Alexa488 (1:200, 53-9760-82, Thermo Fisher Scientific, Massachusetts, USA). Anti-α-smooth muscle actin was added after a series of PBS washes following the first staining. Nuclei were stained with DAPI (1.24653, Merck, Darmstadt, Germany) and slides were mounted with SlowFade Antifade mounting medium (S36937, Thermo Fisher Scientific, Massachusetts, USA).

Images were captured Zeiss (LSM 710, Unidad de Citometría de Flujo y Microscopía de Fluorescencia, Universidad Complutense de Madrid) laser-scanning confocal microscope from each section. For Z stack images, 4-8 consecutive XY images were obtained on the Z axis by Zeiss confocal microscope (LSM 710). ImageJ software was used to analyse the collected images.

### Superoxide measurements by DHE and lucigenin chemiluminescence

OCT-embedded penis sections from humans and rats of 6-8 μm were cut in a cryostat and incubated with or without 4-Hydroxi-TEMPO (4-TEMPOL; Merck, Darmstadt, Germany) 10mM for 30 minutes. Afterwards, rat slices were exposed to 3 μM dihydroethidium (DHE) while in humans’ sections it was added 4 μM DHE (Merck, Darmstadt, Germany) for 30 minutes at 37ºC ^51^. Nuclei were stained with 1 μM DAPI for 5 minutes. All images were taken by fluorescence microscope (Leica microsystems, Wetzlar, Alemania). DHE intensity was obtained through ImageJ software and normalized with DAPI intensity.

CC strips were dissected and then transferred to microtiter plate wells containing 5 μM bis-N-methylacridinium nitrate (Lucigenin; Merck, Darmstadt, Germany), some strips were stimulated with NADPH (100 μM; Merck, Darmstadt, Germany) and 4-TEMPOL (1 mM) was used as negative control. Chemiluminescence was measured in a luminometer (BMG FluostarOptima). Baseline values were subtracted from the counting values under the different experimental conditions and superoxide production was normalized to dry tissue weight.

### Measurement of 25-hydroxyvitamin D and testosterone in plasma

25-hydroxyvitamin D and testosterone levels were measured in rat citrated plasma and human EDTA plasma using a General 25-Hydroxyvitamin D3 (HVD3) ELISA Kit (Reddot Biotech Inc., Kelowna, Canada) and Testosterone ELISA DE1559 (Demeditec, Kiel, Germany) following manufacturer’s instructions.

### Statistics

Analysis was performed using GraphPad Software v8 (GraphPad Software Inc., USA). All data were tested for normal distribution using the Shapiro-Wilk test and parametric or non-parametric statistics were used as appropriate. Data are presented either as scatter plots and means or as means ± SEM. Two-way ANOVA analysis and for multiple comparisons the Sidak method or the Bonferroni post hoc test were used. P values of less than 0.05 were considered statistically significant. Pearson correlations were applied when relevant.

