## Supplementary Table and Figures for "Vitamin D deficiency induces erectile dysfunction: role of superoxide"

**Supplemental Table 1. Characteristics of human subjects.** Data from all subjects as well as segregated data from subjects with plasma vitamin D (vitD) levels below the median (<14.2 ng/ml) or above the median (>14.2 ng/ml) are presented. Numerical variables are expressed as mean±SEM and were compared by unpaired Mann-Whiney U-test while categorical variables were compared by Fisher's exact test.

| Variable | all | vitD <14.2 ng/ml | vitD >14.2 ng/ml | P value |
|---|---|---|---|---|
| **n** | 12 | 6 | 6 | |
| **Age (years)** | 54.9±4.0 | 54.3±6.6 | 55.5±5.2 | 0.7857 |
| **Diabetes (%)** | 4 (33.3) | 2 (33.3) | 2 (33.3) | 1.0000 |
| **Dislipidemia (%)** | 1 (8.3) | 1 (16.7) | 0 (0.0) | 1.0000 |
| **Hypertension (%)** | 4 (33.3) | 2 (33.3) | 2 (33.3) | 1.0000 |
| **CVD (%)** | 2 (16.7) | 1 (16.7) | 1 (16.7) | 1.0000 |
| **Serum creatinine (mg/dl)** | 1.25±0.15 | 1.42±0.32 | 1.12±0.04 | 0.6970 |

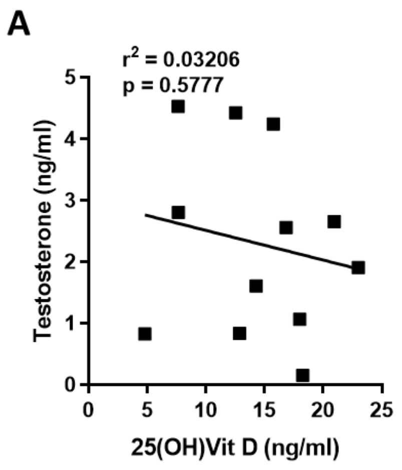


**Supplemental Figure 1.** Vitamin D levels does not correlate with testosterone levels in human plasma donors. Pearson correlation coefficient was calculated.

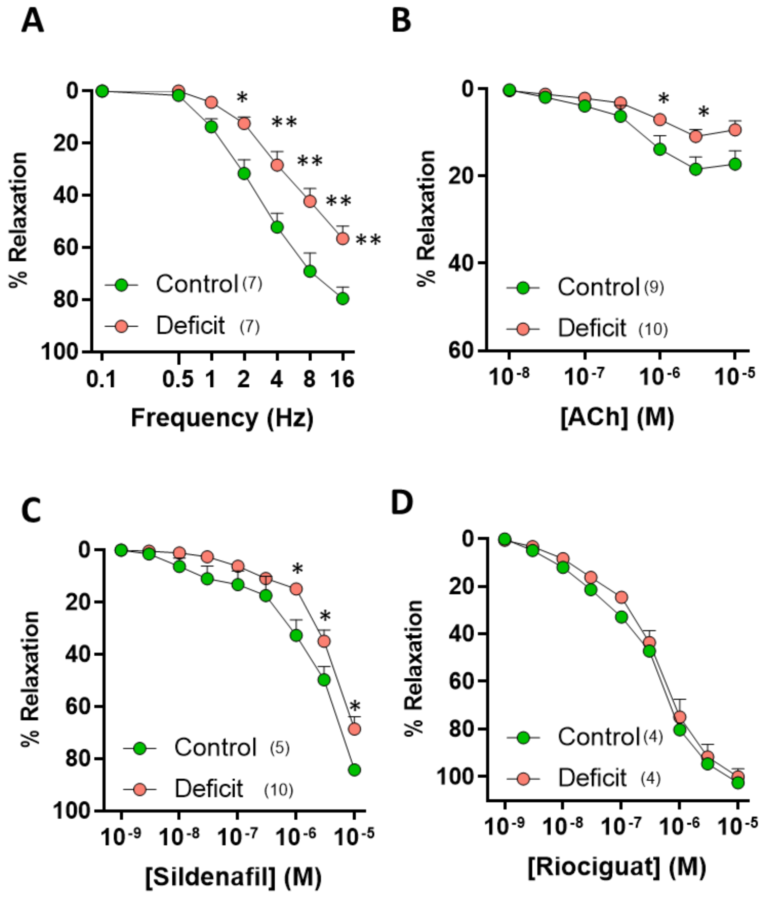


**Supplemental Figure 2. Vitamin D deficiency induces ex vivo erectile dysfunction in male** Sprague Dawley rats. Effect of vitamin D deficit on the relaxant responses in corpora cavernosa induced by EFS (A), acetylcholine (ACh, B), sildenafil (C), and riociguat (D). Results are means ± standard error of the mean. * indicates P<0.05, **P<0.01 vs Control using two-way (deficit x frequency or deficit x concentration) ANOVA test followed by a a Sidak's multiple comparisons test. The number of experiments from different rats is indicated in parenthesis.

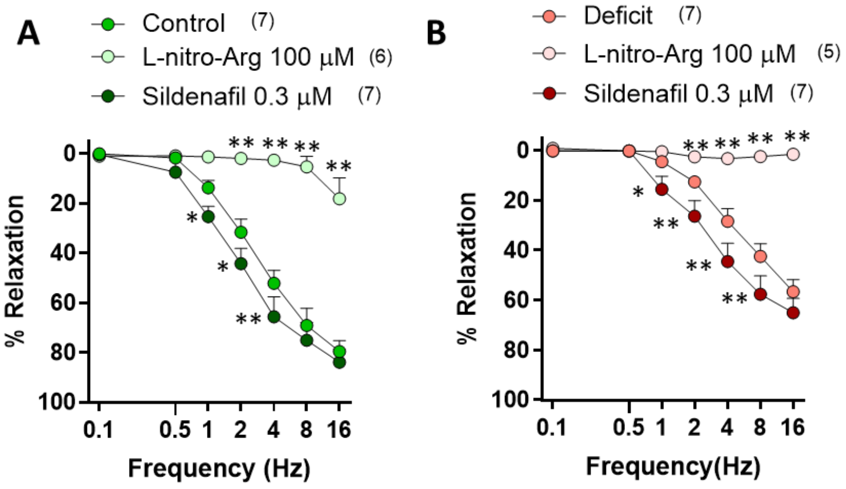


**Supplemental Figure 3. Potentiation by sildenafil and inhibition by L-nitro-Arg of EFS-induced relaxation of CC.** Effect of sildenafil (0.3 μM) and L-nitro-Arg (100 μM) on the curve of relaxation induced by EFS in CC from control (A) and vitamin D deficit (B) Sprague Dawley rats. Results are means ± standard error of the mean. * indicates $P<0.05$, **$P<0.01$ vs Control using two-way (treatment x frequency) ANOVA test followed by a Sidak's multiple comparisons test. The number of experiments from different rats is indicated in parenthesis.

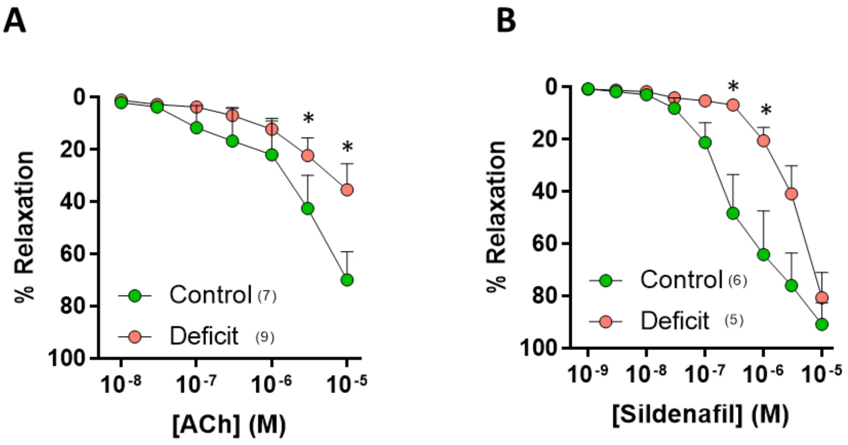


**Supplemental Figure 4. Vitamin D deficiency induces endothelial dysfunction in penile dorsal arteries from male Sprague Dawley (SD) rats.** Effects of vitD deficiency on relaxant responses in penile dorsal arteries induced by A) acetylcholine (ACh) and B) sildenafil. Results are means ± standard error of the mean. * indicates $P<0.05$ using two-way (deficit x concentration) ANOVA test followed by a Bonferroni post hoc test. The number of experiments from different rats is indicated in parenthesis.

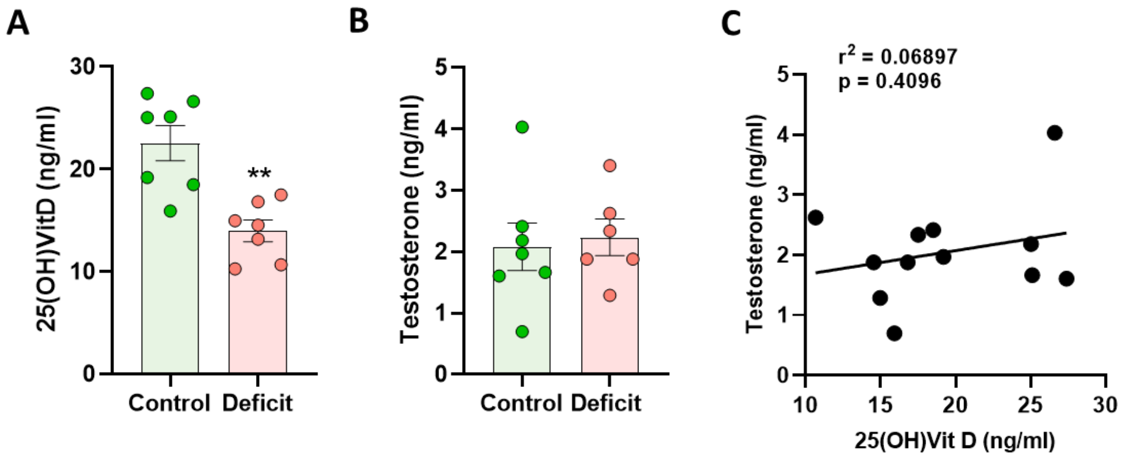


**Supplemental Figure 5. Vitamin D levels does not correlate with testosterone levels in male Wistar rats.** A,B) Effect of a 5-months vitD-free diet in A) 25(OH)VitD plasma levels and B) testosterone plasma levels. C) Pearson correlation was performed between vitD and testosterone plasma level. Results are means ± standard error of the mean. ** indicates P<0.01 using two-way (deficit x concentration) ANOVA test followed by a Bonferroni post hoc test. n=6-7 animals.

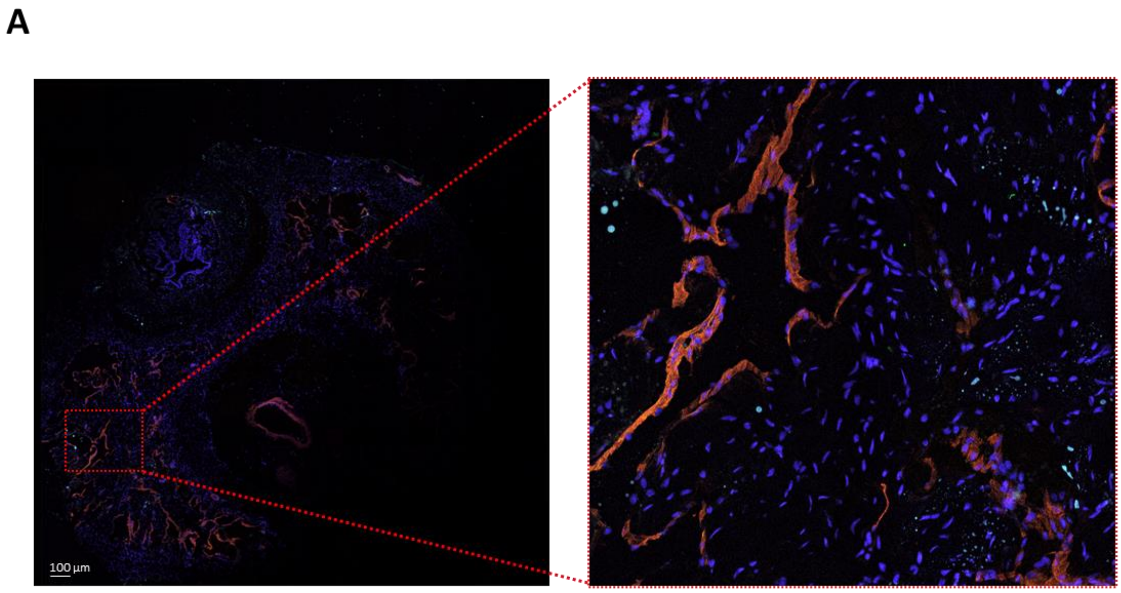


**Supplemental Figure 6.** A) Negative control for VDR immunohistochemistry in the absence of primary antibody anti-VDR, α-actin in red and nuclei are shown in blue (DAPI)
